# A Disease-Agnostic Nasal Microbiome Wellness Index for Standardized Assessment of Upper-Airway Respiratory Health

**DOI:** 10.64898/2026.07.19.739383

**Authors:** Maria A. Barrera-Suarez, Vinod K. Gupta, Xiaowei Zhao, Adam L. Koller, Conan Y. Zhao, Gina A. Suh, Lioudmila V. Karnatovskaia, Erin K. O’Brien, Vanessa L. Kronzer, Jaeyun Sung

## Abstract

**Background:** Upper-airway and respiratory conditions impose a large and growing global burden, yet there is no standardized way to assess whether a nasal microbiome is “healthy”. Currently, monitoring remains reactive, beginning only after symptoms manifest. The nasal cavity is well suited to proactive monitoring: it shapes respiratory health and pathogen colonization resistance, and can be sampled non-invasively and repeatedly. To address this gap, we introduce the Nasal Microbiome Wellness Index (NMWI), a disease-agnostic, continuous score of nasal microbiome health derived from LASSO-penalized logistic regression. Rather than counting taxa, it learns which taxa (and in what balance) characterize a healthy nose and returns the predicted log-odds that a profile resembles a healthy state. The index was trained on 1654 nasal 16S rRNA gene amplicon sequencing samples (589 healthy, 1065 non-healthy) pooled from 27 publicly available studies, uniformly reprocessed through a single computational pipeline.

**Results:** The NMWI comprises an interpretable signature of 24 taxa whose combined relative abundances determine the health-associated log-odds. Health-associated genera such as *Corynebacterium* and *Cutibacterium* raised the score, while dysbiosis-associated genera such as *Pseudomonas* and *Escherichia*–*Shigella* lowered it. The NMWI substantially outperformed the Shannon, Simpson, and Chao1 diversity indices, which showed negligible, directionally inconsistent separation between healthy and non-healthy samples (|Cliff’s *δ*| = 0.01–0.20), whereas the NMWI produced large, consistent separation (*δ* = 0.65). It achieved a balanced accuracy of 74.37% on the training data (resubstitution estimate), and 73.43% under repeated 10-times 10-fold cross-validation. Performance remained stable at mean balanced accuracy of 73.82% under a leave-one-study-out framework, reflecting cross-study generalizability. In independent external cohorts, the balanced accuracy was 71.49%, and leave-one-disease-out analysis (in which each disease condition was withheld from training) showed a mean balanced accuracy of 63.92% across unseen conditions, consistent with a disease-agnostic design. The index also generalized across heterogeneous datasets spanning multiple 16S rRNA gene hypervariable regions—to our knowledge the first demonstration of such cross-study, cross-region transferability for the nasal cavity.

**Conclusions:** The NMWI distills a complex nasal microbial profile into a single interpretable score computed directly from the 16S rRNA gene data that dominate existing nasal research, making it immediately applicable to published and future datasets without re-sequencing. By replacing descriptive, diversity-based comparison with a quantitative standard, it offers a reproducible, open-source foundation for cross-study benchmarking, individual-level phenotyping, and longitudinal respiratory wellness monitoring.

## Background

The human nasal cavity serves as a critical interface between the external environment and the upper-airway respiratory system [1,2]. Unlike the gut, it harbors a relatively low-diversity microbial community dominated by a limited set of commensals and pathobionts, together with viruses and fungi [3,4]. This community varies widely across individuals, shaped by a complex interplay of host, lifestyle, and environmental factors, including age, genetics, immune status, smoking, antibiotic use, and airborne exposures [5–9]. The nasal cavity itself is a high-stress, resource-limited niche characterized by fluctuating humidity, scarce nutrients, and robust host antimicrobial defenses, which collectively restrict colonization to specialized taxa [10,11]. Within this constrained ecosystem, commensal bacteria can modulate host immune responses [12–15], influence colonization resistance [16,17], and limit pathobiont growth [3,18–20]. In particular, commensal–pathobiont interactions are often antagonistic, with several culture-based studies elucidating mechanisms by which commensals inhibit opportunistic growth and thereby shape nasal microbiome composition [16,21–23]. Although pathobionts are typically benign at low abundance, disruptions in community structure can promote pathogenic behavior [24,25]. Consequently, such perturbations may shift host–microbe interactions in ways that favor pathogen expansion, impaired colonization resistance, and disease progression.

The nasal microbiome also has important and clinically consequential links to host health [3–5]. For example, nasal dysbiosis has been linked to diverse disease states and host responses, including local and systemic inflammatory disorders [26–39]; neurodegenerative processes [40–43]; differential antiviral immune responses [44]; susceptibility to respiratory infection [12,45,46]; variability in corticosteroid treatment response [47]; increased disease severity and exacerbations [31,48–50]; and increased risk of lung transplant rejection [51,52]—associations that together impose a substantial burden on healthcare systems worldwide [53–56]. These findings position the nasal microbiome as a compelling target for precision medicine, one that could capture individualized microbial signatures of health and disease and motivate nasal microbiome-targeted strategies for prevention and treatment [57,58]. Critically, the nasal cavity is particularly well suited for such translational applications because its microbial communities can be sampled non-invasively and repeatedly, enabling longitudinal monitoring, scalable population-level surveillance, and early disease risk stratification [59–61]. Yet current clinical practice does not exploit this potential: upper-airway conditions are typically identified and managed only after symptoms become overt (through clinical evaluation, imaging, endoscopy, or escalating therapy) with no scalable, non-invasive means of monitoring nasal microbiome health longitudinally before clinical deterioration occurs. In this regard, a quantitative, repeatable measure of microbiome health could support earlier risk stratification and more proactive management.

To date, most nasal microbiome studies have relied on conventional analytical approaches, such as α-diversity metrics and differential abundance testing, to characterize microbiome differences across health and disease. While informative at a group level, these approaches have well-documented limitations: α-diversity estimates, in particular, are sensitive to sequencing depth, susceptible to incomplete sampling bias, and subject to unaccounted measurement error, all of which can obscure true biological signals and produce findings that do not replicate across datasets [62–66]. More fundamentally, diversity metrics summarize communities by richness and evenness without accounting for taxon identity or the directionality of community shifts. As a result, similar diversity values may correspond to biologically distinct states—for example, comparable diversity scores may arise whether driven by a loss of key commensals or by an expansion of pathobionts, yet these scenarios carry entirely different health implications. This is a critical blind spot in the nasal context, where communities are typically dominated by a few taxa and can transition between distinct low-diversity, taxon-dominated configurations [67]. Indeed, respiratory tract diseases have been associated with altered nasal microbial diversity, yet the direction of these changes is inconsistent across the literature [4,68–70]. These limitations collectively highlight the need for a standardized, interpretable metric that incorporates taxon identity and captures biologically meaningful community shifts, while remaining disease-agnostic, robust to cross-study technical heterogeneity, and actionable at the individual-patient level.

In this study, we introduce the Nasal Microbiome Wellness Index (NMWI), a standardized, disease-agnostic metric that evaluates nasal microbiome health by estimating the likelihood that an individual’s microbial community resembles configurations observed in healthy individuals. The NMWI is conceptually inspired by the Gut Microbiome Wellness Index (GMWI) [71] and its refinement GMWI2 [72], which distill complex gut microbiome profiles into a single interpretable health score. Extending this framework to the nasal cavity, we derived the NMWI using a LASSO-penalized logistic regression classifier trained on 1654 nasal microbiome profiles, including 589 healthy and 1065 non-healthy samples, retrieved from 27 publicly available studies spanning seven chronic disease conditions across four continents. The resulting score—defined as the predicted log-odds of belonging to a healthy microbiome state—integrates signals across multiple microbial taxa into a single continuous value, providing a more ecologically informed and generalizable summary of nasal microbiome health than any individual taxon or diversity metric alone. Here, we demonstrate that the NMWI achieves robust cross-study and cross-disease generalizability, outperforms conventional α-diversity metrics, and establishes an open-source, extensible tool for nasal microbiome-based phenotyping and precision respiratory medicine.

## Methods

### Study identification and eligibility criteria

A systematic search of PubMed and the NCBI Sequence Read Archive (SRA) was conducted using the keywords “nasal microbiome”, “16S rRNA”, and “nasal cavity” to identify publicly available studies profiling the adult human nasal microbiome. The literature search covered publications through August 2025. Eligibility was restricted to studies sampling the nasal cavity or sinonasal regions. As previously defined [73], the nasal cavity extends from the nostrils (nares) to the choanae; studies sampling the nasopharynx or lower respiratory tract were therefore excluded. When anatomical landmarks were not explicitly reported, sampling depth (1–7 cm from the nares) was used as a proxy for nasal cavity sampling. Samples from the ethmoid and maxillary sinuses were retained given the well-documented compositional overlap among sinonasal microbial communities [11,74,75].

Throughout this study, the unit of analysis is the individual nasal microbiome sample, i.e., a single specimen collected from one participant at one anatomical site and time point by nasal swabbing, nasal lavage, or nasal brushing. The definitions of “healthy” and “non-healthy” are consistent with our prior work on gut microbiome wellness indices. Specifically, “healthy” refers to individuals with no reported diagnosis of a disease or disease-related condition at the time of sampling. “Non-healthy” refers to individuals with a confirmed clinical diagnosis of a chronic disease, sampled during a stable disease interval to avoid the transient, condition-specific perturbations characteristic of acute illness [31,49]. These definitions are intentionally disease-agnostic: the non-healthy group encompasses patients with diverse chronic conditions rather than any single diagnosis, reflecting our goal of identifying generalizable microbial signatures of health status rather than disease-specific signatures.

Eligible datasets (**Additional file 1: Tables S1, S2**) met all of the following criteria: (i) association with a peer-reviewed publication; (ii) 16S rRNA gene sequencing performed on an Illumina platform; (iii) inclusion of adult participants (≥18 years) to minimize age-related compositional effects [6]; (iv) public availability of raw FASTQ files with accompanying sample-level metadata; and (v) representation of each chronic disease condition by at least two independent studies to mitigate single-cohort bias. Samples annotated for current antibiotic use or active smoking were excluded *a priori*. For participants whose samples were collected longitudinally, only the earliest time point was retained.

### 16S rRNA gene amplicon sequence data processing

Raw FASTQ files and associated metadata were downloaded from the NCBI SRA. Study-level metadata, including the targeted 16S rRNA gene hypervariable region, anatomical sampling location, geographic origin of the cohort, clinical phenotype, sequencing platform, and sample collection method, were curated from each study’s main text, supplementary materials, and associated public repositories. Some host physiological variables (e.g., sex, age, BMI, and disease-specific measurements) were not consistently reported across studies and were therefore excluded from further analyses.

To ensure comparability across heterogeneous datasets and to minimize biases arising from variable reverse-read quality and merging efficiency, only forward reads were retained for processing. All sequence data were processed through a single, uniformly applied bioinformatics pipeline: Sequences were processed in QIIME 2 version 2024.10.1 [76] using the DADA2 plugin [77] for quality filtering, denoising, chimera removal, and inference of amplicon sequence variants (ASVs). Reads were trimmed 15 bp from the 5′ end (--p-trim-left 15) and truncated at the first position where the Phred quality score dropped to Q ≤ 20. All other DADA2 parameters were left at their default values, which were deemed appropriate following visual inspection of quality profiles across representative samples from multiple studies.

Rarefaction curves identified a minimum sequencing depth of 9231 reads, below which samples were excluded to ensure sufficient coverage for downstream diversity analyses. Two additional filters were then applied: individual samples containing fewer than five ASVs were removed, and any study with a final sample size below 10 was excluded. This standardized pipeline was applied uniformly to all datasets despite differences in targeted 16S rRNA hypervariable regions, enabling direct cross-study comparability while minimizing technical batch effects.

### Taxonomic annotation of ASVs

ASVs were assigned taxonomy using a Naïve Bayes classifier trained on the SILVA full-length 16S rRNA gene database v138 [78], as previously described [68]. Features lacking phylum-level classification or assigned to chloroplasts, mitochondria, archaea, or eukaryota were removed from the ASV feature table. The table was then collapsed to the genus level for downstream analyses, with higher taxonomic ranks (family, order, class, and phylum) retained for ASVs that could not be resolved to genus. Finally, to reduce the influence of sparsely observed taxa, a prevalence filter was applied to remove features present in fewer than 5% of samples in the pooled dataset.

### Statistical and bioinformatic analyses

All downstream analyses were performed in Python v3.12.2 [79] unless otherwise specified. α-diversity was quantified at the genus level on rarefied feature tables using the Shannon, Simpson, and Chao1 indices to capture complementary aspects of community evenness, dominance, and richness. β-diversity was assessed using Bray–Curtis dissimilarity computed on genus-level relative abundances. Differences in community composition between healthy and non-healthy groups were tested using permutational multivariate analysis of variance (PERMANOVA) implemented via the adonis2 function in the R package vegan [80,81], with 999 permutations stratified by study to account for study-specific effects. Differential abundance between healthy and non-healthy groups was assessed at the genus level using Analysis of Compositions of Microbiomes with Bias Correction 2 (ANCOM-BC2) [82], which corrects for compositional bias and sampling fraction differences across samples. Specifically, health status was included as a fixed effect with healthy samples as the reference group, and structural zeros were incorporated into the model. Multiple hypotheses were controlled using Holm-adjusted *P*-values, with adjusted *P*-values < 0.05 and |natural log-fold change | > 0.5 considered significant. Positive ln(fold change) indicated higher abundance in non-healthy samples, whereas negative values indicated higher abundance in healthy samples.

### Construction of the Nasal Microbiome Wellness Index (NMWI)

Following the modeling framework of GMWI2 [72], we constructed the NMWI as an L1-penalized (LASSO) logistic regression classifier that distinguishes healthy from non-healthy nasal microbiome profiles. This formulation imposes an L1 penalty on the model coefficients, controlled by the regularization parameter *C* (where smaller values of *C* induce stronger shrinkage) [83]. The penalty encourages sparsity by driving coefficients of less informative taxa toward zero, thereby selecting a parsimonious subset of predictive microbial features. The response variable encodes health status, while the predictor matrix comprised genus-level relative abundances where available; when an ASV could not be confidently assigned to a particular genus, the highest confidently resolved taxonomic rank (e.g., family or order) was retained and included as a predictor.

Importantly, the NMWI is fundamentally a continuous score: positive values indicate greater resemblance to healthy microbiome configurations, negative values indicate a shift toward non-healthy states, and the absolute magnitude reflects the model’s confidence in the prediction. Binary classification is an optional downstream step that can be applied by thresholding the score at zero (or at a user-defined magnitude cutoff) but the score itself carries graded, quantitative information that is lost under forced binary assignment. This design mirrors the original GMWI2 framework [72], in which the predicted log-odds score is likewise treated as a continuous health indicator rather than a classifier output.

Hyperparameter optimization was performed via grid search over a range of candidate *C* values within a leave-one-study-out cross-validation (LOSO CV) strategy. In each iteration, all samples from one entire study were held out as an independent test set while the remaining studies were used for training. The LASSO model was implemented in scikit-learn v1.7.2 [84] using the LIBLINEAR solver [85] with class weights set to “balanced” to address label imbalance. Model performance at each *C* value was evaluated by balanced accuracy (the mean of sensitivity and specificity) averaged across held-out studies. The optimal *C* was selected to maximize this metric. A detailed mathematical description of the LASSO-penalized logistic regression objective, coefficient learning, and score computation is provided in the GMWI2 study. The NMWI implementation follows the identical rationale and software infrastructure. Complete hyperparameter grids, cross-validation scripts, and trained model weights are available in the project repository (https://github.com/mabarrera12/NMWI/).

### Model training and performance evaluation

Classification performance was assessed using the LOSO CV framework described above, as well as 10-times repeated 10-fold cross-validation (10×10 CV) on the pooled dataset. After hyperparameter tuning, the final L1-regularized logistic regression model was fit on the entire training cohort (1654 samples) at the optimal *C*. The resulting NMWI is defined as the predicted log-odds of belonging to the healthy class, computed as a sparse linear combination of genus-level (and higher-rank) relative abundances weighted by the learned coefficients. This yields a compact, interpretable representation of nasal microbiome health.

Performance was quantified by balanced accuracy and the area under the receiver operating characteristic curve (AUROC). To provide a stringent test of generalizability beyond the discovery data, the model was prospectively evaluated on completely independent external cohorts and disease conditions that were not part of the training dataset (**Additional file 1: Tables S3, S4**).

### Magnitude-based classification using NMWI

Because the sign of the NMWI indicates predicted health status (positive, healthy-like; negative, non-healthy-like) and its absolute magnitude reflects prediction confidence, discrete classification can be made more conservative by applying a symmetric magnitude cutoff *t*: samples with |NMWI| ≥ *t* are assigned to a class, while samples below the decision boundary (|NMWI| < *t*) are left indeterminate. Classification performance across a range of *t* values was evaluated using balanced accuracy and AUROC within both the 10×10 CV and resubstitution frameworks. This thresholding approach explicitly trades off coverage for higher predictive certainty, mirroring common practices in clinical ML deployment where high-confidence decisions are prioritized over forced calls.

## Results

### Study selection and characteristics of the included cohorts

To construct the NMWI, we systematically assembled publicly available nasal microbiome datasets from PubMed and the NCBI Sequence Read Archive (see **Methods**). Eligible studies provided 16S rRNA gene sequencing data together with metadata on anatomical sampling site, clinical phenotype, sequencing platform, targeted hypervariable region, and demographic characteristics. To ensure comparability across studies and minimize technical variability introduced by heterogeneous analytical workflows, all sequencing data were reprocessed through a single standardized bioinformatics pipeline. The full workflow, from dataset identification through final sample retention, is described in **Figure 1a**.

After data acquisition, quality control, and application of the selection criteria, 1654 samples were retained for model construction, comprising 589 healthy and 1065 non-healthy samples (**Table 1**; **Additional file 1: Tables S1, S2**). These samples derived from 27 studies spanning four continents (**Fig. 1b**) and the following seven chronic disease categories (**Fig. 1c**): allergic rhinitis (AR), asthma (AS), chronic rhinosinusitis (CRS), CRS with comorbid asthma (CRS-AS), cystic fibrosis (CF), granulomatosis with polyangiitis (GPA), and rheumatoid arthritis (RA). A separate, independent validation set of 350 samples (109 healthy, 241 non-healthy) from 11 independent studies was reserved for external evaluation (**Table 1**; **Additional file 1: Tables S3, S4**).

**Table 1.** Characteristics of all studies included in the index and validation datasets.

| First author and publication year | Clinical phenotype | Sequencing platform | Sampling location | Country of origin | 16S rRNA gene region | NCBI BioProject accession | No. of samples in index training (Validation) | Ref. |
| --- | --- | --- | --- | --- | --- | --- | --- | --- |
| Wang *et al.*, 2023 | AR | Illumina NovaSeq 6000 | IT | China | V3–V4 | PRJNA899122 | 77 (0) | [86] |
| Mariani *et al.*, 2021 | AR | Illumina MiSeq | AN | Italy | V3–V4 | PRJNA646474 | 43 (0) | [87] |
| Perez-Losada *et al.*, 2023 | AS | Illumina MiSeq | AN | Portugal | V4 | PRJNA913468 | 31 (0) | [69] |
| Durack *et al.*, 2018 | AS | Illumina MiSeq | MM | USA | V4 | PRJEB22676 | 28 (0) | [88] |
| Michalovich *et al.*, 2019 | AS | Illumina MiSeq | MM | Poland | V4 | PRJNA434133 | 17 (0) | [89] |
| Chen *et al.*, 2022 | AS, AR | Illumina HiSeq 2500 | MM | China | V3–V4 | PRJNA793600 | 87 (0) | [90] |
| Armbruster *et al.*, 2024 | CF | Illumina NovaSeq 6000 | Sinus | USA | V4 | PRJNA1081394 | 16 (0) | [91] |
| Voronina *et al.*, 2020 | CF | Illumina MiSeq | Sinus | Russia | V1–V4 | PRJNA544655 | 14 (0) | [92] |
| Boeck *et al.*, 2017 | CRS, CRS-AS | Illumina MiSeq | AN | Belgium | V4 | PRJEB30316 | 185 (0) | [74] |
| Wagner *et al.*, 2019 | CRS | Illumina MiSeq | MM | New Zealand | V4 | PRJNA512906 | 24 (0) | [93] |
| Cope *et al.*, 2017 | CRS | Illumina MiSeq | Sinus | USA | V4 | PRJEB20232 | 17 (0) | [94] |
| Lucas *et al.*, 2021 | CRS-AS | Illumina MiSeq | Sinus | USA | V4 | PRJNA656590 | 11 (0) | [95] |
| Liang *et al.*, 2025 | CRS | Illumina HiSeq 2500 | MM | China | V3–V4 | PRJNA1028156 | 160 (0) | [96] |
| Szaleniec *et al.*, 2024 | CRS | Illumina MiSeq | MM | Poland | V3–V4 | PRJEB55924 | 142 (0) | [97] |
| Hoggard *et al.*, 2017 | CRS | Illumina MiSeq | MM | New Zealand | V3–V4 | PRJNA338893 | 131 (0) | [98] |
| Liang *et al.*, 2023 | CRS | Illumina HiSeq 2500 | MM | China | V3–V4 | PRJNA785109 | 115 (0) | [34] |
| Bartosik *et al.*, 2023 | CRS | Illumina MiSeq | MM | Austria | V3–V4 | PRJNA907202 | 31 (0) | [99] |
| Koeller *et al.*, 2018 | CRS | Illumina MiSeq | MM | USA | V3–V4 | PRJEB23675 | 23 (0) | [75] |
| Biswas *et al.*, 2017 | CRS | Illumina MiSeq | MM | New Zealand | V3–V4 | PRJNA390854 | 13 (0) | [100] |
| Gan *et al.*, 2020 | CRS, AR | Illumina MiSeq | MM | China | V3–V4 | PRJNA493980 | 113 (0) | [101] |
| Shi *et al.*, 2021 | CRS, NK/T | Illumina HiSeq 2500 | AN | China | V4 | PRJNA733573 | 32 (27) | [102] |
| Rhee *et al.*, 2018 | GPA | Illumina MiSeq | MM | USA | V1–V2 | PRJNA473785 | 97 (0) | [103] |
| Lamprecht *et al.*, 2019 | GPA, RA | Illumina MiSeq | AN | Germany | V3–V4 | PRJNA494384 | 33 (0) | [104] |
| Koskienen *et al.*, 2018 | Healthy | Illumina MiSeq | OC | Austria | V4 | PRJEB20526 | 52 (0) | [105] |
| Cleary *et al.*, 2021 | Healthy | Illumina NextSeq 500 | AN | Malaysia | V4 | PRJEB38610 | 12 (0) | [106] |
| van Dalen *et al.*, 2023 | Healthy | Illumina MiSeq | AN | Germany | V1–V3 | PRJNA923578 | 122 (0) | [107] |
| Bin *et al.*, 2024 | RA | Illumina MiSeq | MM | South Korea | V3–V4 | PRJNA1058141 | 28 (0) | [108] |
| van den Munckhof *et al.*, 2020 | RTI | Illumina MiSeq | AN | Netherlands | V3–V4 | PRJNA596902 | 0 (69) | [109] |
| Heikema *et al.*, 2020 | RTI | Illumina MiSeq | AN | Netherlands/ Israel | V5–V6 | PRJEB28612 | 0 (37) | [110] |
| Fernandez-Rodriguez *et al.*, 2024 | Healthy | Illumina MiSeq | AN | USA | V1–V2 | PRJNA1024520 | 0 (19) | [111] |
| Hickman *et al.*, 2024 | Healthy | Illumina MiSeq | MM | USA | V3–V4 | PRJNA746950 | 0 (18) | [112] |
| Ryser *et al.*, 2025 | Healthy | Illumina NovaSeq 6000 | AN | Switzerland | V3–V4 | PRJNA1213741 | 0 (22) | [113] |
| Nardelli *et al.*, 2022 | COVID-19 | Illumina MiSeq | AN | Italy | V1–V3 | PRJNA744167 | 0 (15) | [114] |
| Rhoades *et al.*, 2021 | COVID-19 | Illumina MiSeq | AN | USA | V4 | PRJNA745169 | 0 (73) | [115] |
| Wu *et al.*, 2023 | OS | Illumina MiSeq | Sinus / MM | China | V3–V4 | PRJNA930496 | 0 (59) | [116] |
| Garcia-Núñez *et al.*, 2020 | CF | Illumina MiSeq | MM | Spain | V3–V4 | PRJEB31332 | 0 (4) | [117] |
| Lucas *et al.*, 2018 | CF | Illumina MiSeq | MM | USA | V4 | PRJNA374847 | 0 (7) | [118] |

Abbreviations: AN, anterior nares; AR, allergic rhinitis; AS, asthma; CF, cystic fibrosis; CRS, chronic rhinosinusitis; CRS-AS, CRS with comorbid asthma; GPA, granulomatosis with polyangiitis; IT, inferior turbinate; MM, middle meatus; NK/T, NK/T-cell lymphoma; OC, olfactory cleft; OS, odontogenic sinusitis; RA, rheumatoid arthritis; RTI, respiratory tract infection.

### Taxonomic composition and community structure of the nasal microbiome

Following sequence quality control and taxonomic assignment, the resulting feature table was collapsed to the genus level, as species-level classification from 16S rRNA gene data is often unreliable [119,120]. ASVs lacking genus-level annotation were not discarded; instead, taxa resolved only to higher ranks were retained to preserve all measurable taxonomic information. Applying a 5% prevalence threshold to reduce sparsity yielded 184 taxa (comprising 141 genera, 25 families, 10 orders, 6 classes, and 2 phyla) for downstream analysis.

The taxonomic composition of the pooled dataset at the family and genus levels is shown for healthy and non-healthy samples in **Figure 1d**. At the family level, *Corynebacteriaceae*, *Staphylococcaceae*, and *Enterobacteriaceae* were most abundant, together accounting for 47.0% of the total mean relative abundance. At the genus level, communities were dominated by a limited set of genera—primarily *Corynebacterium*, *Staphylococcus*, *Escherichia–Shigella*, and *Cutibacterium*—which collectively accounted for 48.9% of the total mean relative abundance, consistent with prior characterizations of the adult nasal microbiome [37,58,67]. These genera comprised a slightly greater share of the community in healthy (53.7%) than in non-healthy samples (46.2%).

Differential abundance analysis (**Additional file 1: Tables S5, S6**; **Additional file 2: Figures S1, S2**) revealed distinct taxonomic signatures between groups. For example, *Escherichia–Shigella*, *Pseudomonas*, and three others were more abundant in non-healthy samples, whereas *Corynebacterium*, *Cutibacterium*, and six others had higher abundance in healthy samples (ANCOM-BC2, Holm-adjusted *P* < 0.05; |ln(fold change)| > 0.5), in agreement with previous reports [58,75,121]. Notably, *Staphylococcus* abundance did not differ significantly between groups. This genus encompasses species with divergent ecological roles—including *S. aureus* [20,122], *S. epidermidis* [123]*, S. lugdunensis* [23] or *S. hominis* [124]—that the limited resolution of 16S rRNA gene sequencing cannot distinguish, potentially masking biologically meaningful intrageneric shifts.

Beyond relative abundance, we examined taxon prevalence across samples. Before prevalence filtering, 2103 taxa were detected, and their distribution across the 1654 samples was strongly right-skewed (**Fig. 1e**): 1491 taxa (≤ 1% prevalence) were rare, 184 were present in at least 5% of samples, and only eight were detected in ≥ 50%. These eight (*Staphylococcus* (89.0%), *Corynebacterium* (84.9%), *Cutibacterium* (71.5%), *Streptococcus* (66.2%), *Anaerococcus* (63.4%), *Peptoniphilus* (58.4%), *Pseudomonas* (54.4%) and *Acinetobacter* (52.7%)) were the most prevalent in both groups, though their relative proportions differed (**Fig. 1f**).

Finally, to examine differences in microbial community structure, we performed principal coordinates analysis (PCoA) based on Bray–Curtis dissimilarity of genus-level relative abundances (**Fig. 1g**). The analysis revealed a significant difference between healthy and non-healthy samples (PERMANOVA, *R*^2^ = 0.024, *P* < 0.001), even after accounting for study-specific effects by constraining permutations within each study. Although the effect size was modest, these results indicate a consistent shift in nasal microbial community composition associated with disease across independent cohorts.

**Figure 1.**
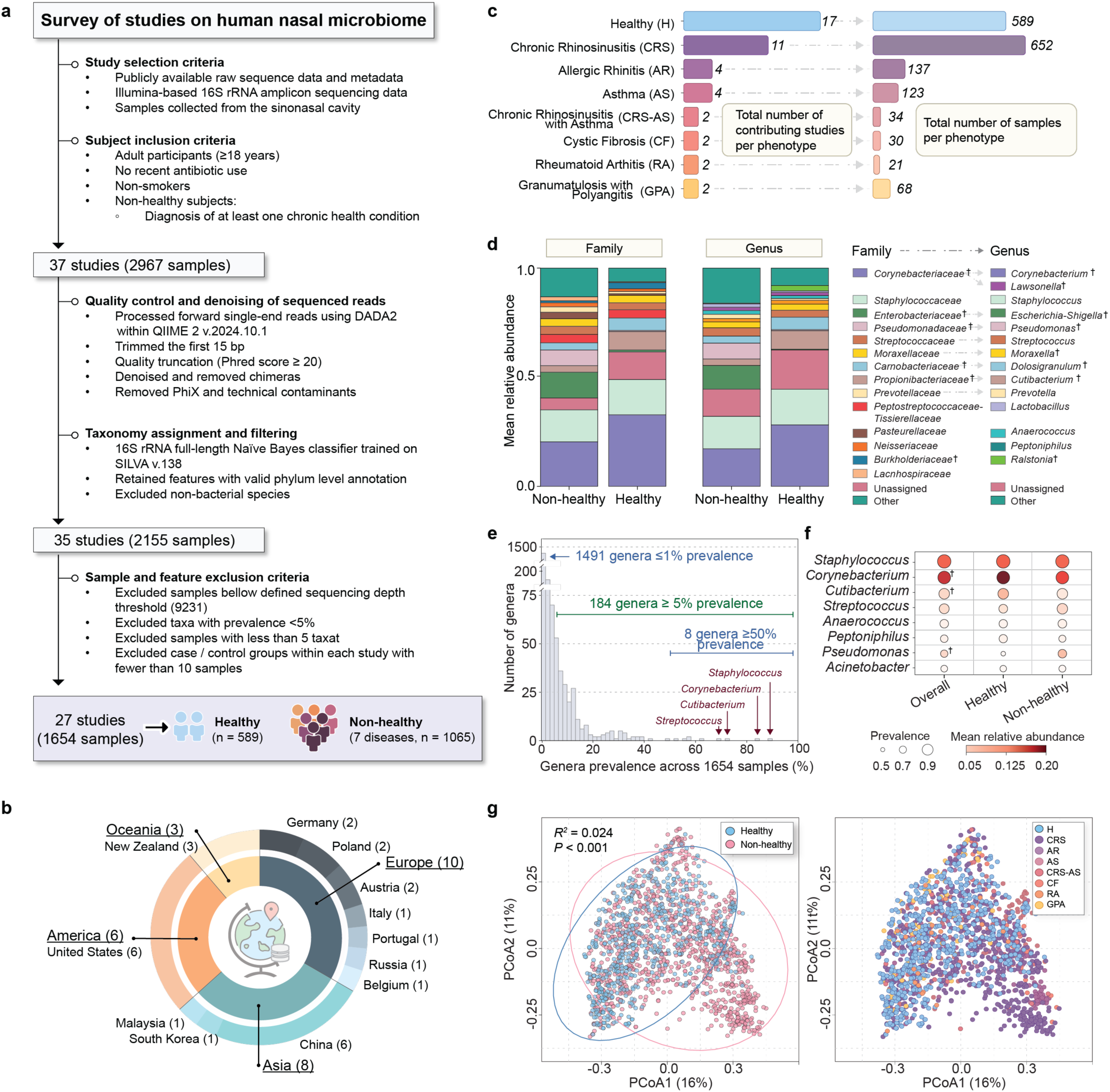
Study selection workflow and characteristics of the data included in NMWI construction. **(a)** Systematic search and filtering: 37 Illumina-based 16S rRNA gene nasal microbiome studies (2967 samples) were identified, of which 27 studies (1654 samples) were retained after preprocessing and filtering. From these 27 studies and 1654 samples: **(b)** Geographic origins across four continents and 12 countries. Numbers in parentheses indicate study counts; **(c)** Number of studies and their samples across seven disease phenotypes and a healthy reference group; **(d)** Mean relative taxonomic composition at the family and genus levels in healthy and non-healthy samples. Arrows link families to representative genera, and crosses (†) denote significantly different taxa (ANCOM-BC2, Holm-adjusted *P* < 0.05); **(e)** Distribution of genus prevalence across all samples; **(f)** Top eight most prevalent genera, with bubble size indicating prevalence and color indicating mean relative abundance (%) per group. Crosses (†) denote significantly different taxa. **(g)** PCoA of Bray–Curtis dissimilarity (PERMANOVA, *R*^2^ = 0.024, *P* < 0.001), with the same ordination colored by health status (left) and disease phenotype (right). Abbreviations: AR, allergic rhinitis; AS, asthma; CF, cystic fibrosis; CRS, chronic rhinosinusitis; CRS-AS, CRS with comorbid asthma; GPA, granulomatosis with polyangiitis; H, healthy; RA, rheumatoid arthritis.

### Construction of the Nasal Microbiome Wellness Index (NMWI)

Following the rationale of GMWI [71] and GMWI2 [72], we modeled the log-odds that a nasal microbiome profile reflects a healthy vs. non-healthy state from genus-level relative abundances. The NMWI was constructed using L1-regularized (LASSO) logistic regression, which simultaneously estimates health status and selects informative microbial predictors. Hyperparameters were tuned within the leave-one-study-out cross-validation (LOSO CV) framework to maximize mean balanced accuracy (see **Methods**), yielding an optimal regularization parameter of *C* = 0.9375 (**Additional file 2: Figure S3**); this strategy improves cross-study performance while preventing within-study data leakage, providing a realistic estimate of generalizability to independent datasets. The final model was trained on the full set of 1654 samples (589 healthy, 1065 non-healthy).

### Microbial features and stability underlying the NMWI

The final model retained 24 taxa (14 with positive and 10 with negative coefficients) that collectively define the NMWI (**Additional file 1: Table S7**). Taxa with positive coefficients raised the NMWI score and were associated with the healthy state, whereas those with negative coefficients lowered the score and were associated with non-healthy states (**Fig. 2a**). Positive-coefficient taxa included *Corynebacteriales, Corynebacteriaceae, Cutibacterium, Ralstonia, Bacteroidia,* and *Actinobacteria*, many of which have been previously linked to a health-associated nasal microbiome [37,58,67]. Conversely, negative-coefficient taxa included *Lachnospiraceae, Pseudomonas*, *Escherichia–Shigella*, *Haemophilus*, *Anaerococcus*, and *Achromobacter* consistent with their reported higher abundance in dysbiotic nasal states [58,69,98,121,125].

To evaluate robustness, we quantified how frequently each taxon was retained across LOSO CV iterations. As shown in **Figure 2a**, bubble size reflects selection frequency, providing a measure of stability. **Figure 2b** presents a heatmap of the logistic regression coefficients across LOSO CV iterations, where each column represents a model trained with one study held out. Importantly, highly stable features exhibited consistent coefficient signs across iterations, indicating robust and reproducible associations with health status. Although coefficient magnitudes varied somewhat, the direction and relative importance of the key taxa were largely preserved, with retention frequencies spanning approximately 53% to 100% across iterations.

**Figure 2.**
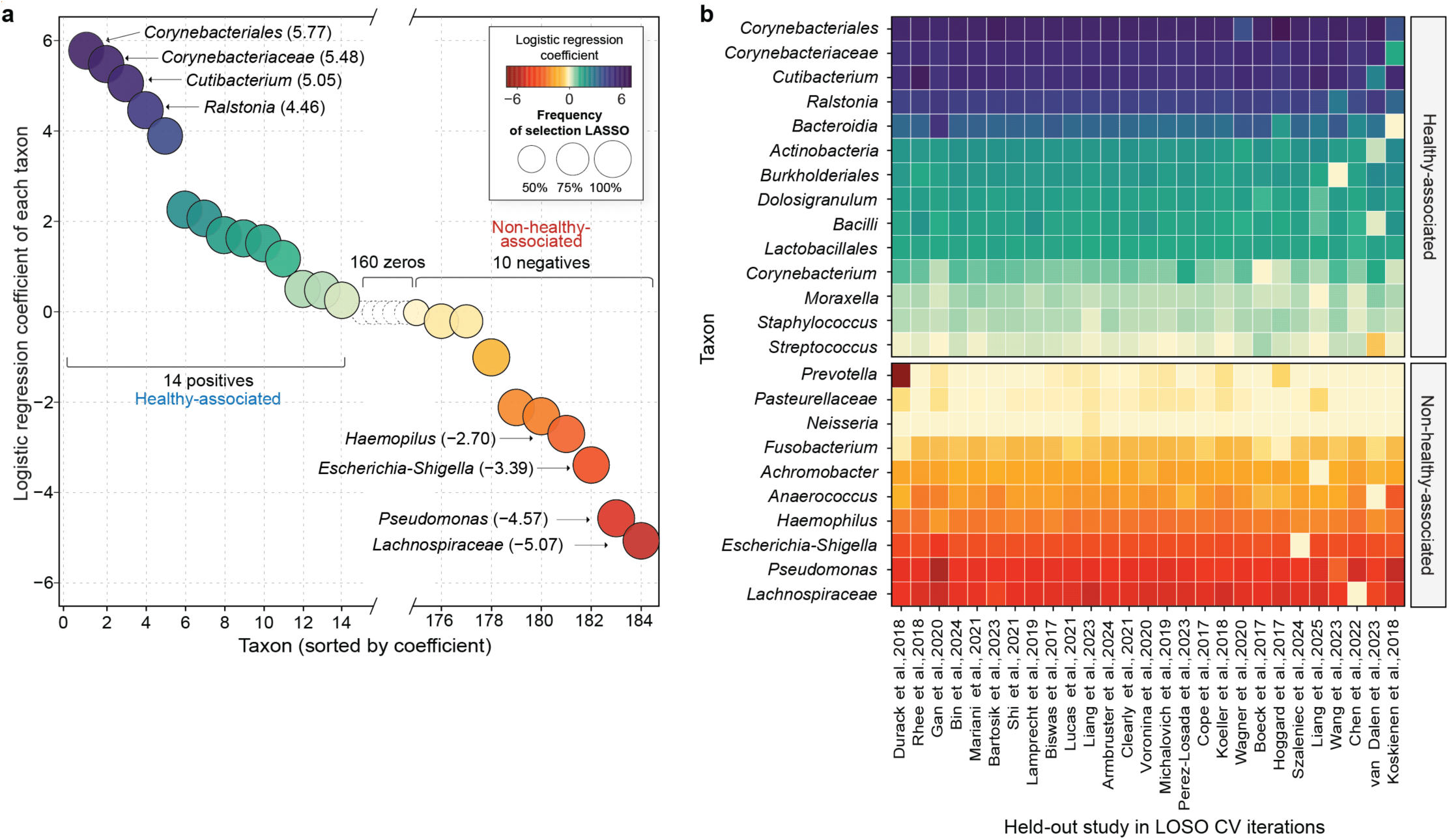
Feature selection and coefficients underlying the NMWI. **(a)** LASSO-penalized logistic regression coefficients: 14 taxa with positive coefficients, 10 with negative coefficients, and 160 with coefficients shrunk to zero. Bubble size indicates selection frequency across leave-one-study-out cross-validation (LOSO CV), and color indicates coefficient magnitude. **(b)** Heatmap of coefficients across LOSO CV models, in which each column represents a model trained with one study held out; and each row a taxon in the final model. Coefficients are represented using the same color scale as in panel **(a)**.

### NMWI outperforms conventional diversity metrics in distinguishing health from disease

We next benchmarked the NMWI against three widely used diversity metrics: Shannon diversity, Simpson index, and Chao1 index. Chao1 richness was significantly higher in non-healthy samples (Mann–Whitney *U* test, *P* = 1.38 × 10^−11^), although the effect size was small (|Cliff’s *δ*|= 0.20). In contrast, Shannon diversity and Simpson index did not differ significantly between healthy and non-healthy samples and exhibited negligible effect sizes (|Cliff’s *δ*| = 0.01 and 0.03, respectively) (**Fig. 3a**). In contrast, NMWI values were significantly higher in healthy compared to non-healthy samples (Mann–Whitney *U* test, *P* = 1.15 × 10^−107^) with a large effect size (|Cliff’s *δ*|= 0.65) and substantially less overlap between groups. The diversity metrics were also directionally inconsistent: the median Chao1 index was slightly higher in non-healthy samples, whereas median Shannon and Simpson values were modestly higher in healthy samples.

To assess disease-specific patterns, we next compared each disease cohort with the pooled healthy population (**Fig. 3b**). NMWI was lower in most disease cohorts than in healthy controls, although the magnitude of separation varied by condition, with effect sizes ranging from 0.12 to 0.87. GPA was the sole exception, showing no significant difference from healthy controls and a slightly positive median NMWI.

Diversity metrics, in contrast, did not exhibit consistent patterns across diseases. Shannon diversity, for example, varied in direction across conditions, with some diseases showing higher median values and others showing lower values relative to healthy samples (**Fig. 3c**). Notably, RA, CRS–AS, and AR samples did not differ significantly from healthy controls, showing substantial distribution overlap (|Cliff’s *δ*| ≤ 0.16). Chao1 behaved similarly (**Additional file 2: Figure S5**; **Additional file 1: Table S8**), varying in direction with no consistent trend (|Cliff’s *δ*| = 0.07–0.67) and failing to reach significance for GPA. The two metrics also differed in the direction of the observed shifts. More specifically, CRS and AR exhibited positive effect sizes for Chao1 but negative effect sizes for Shannon diversity—underscoring the interpretive instability of diversity-based comparisons in this body site.

**Figure 3.**
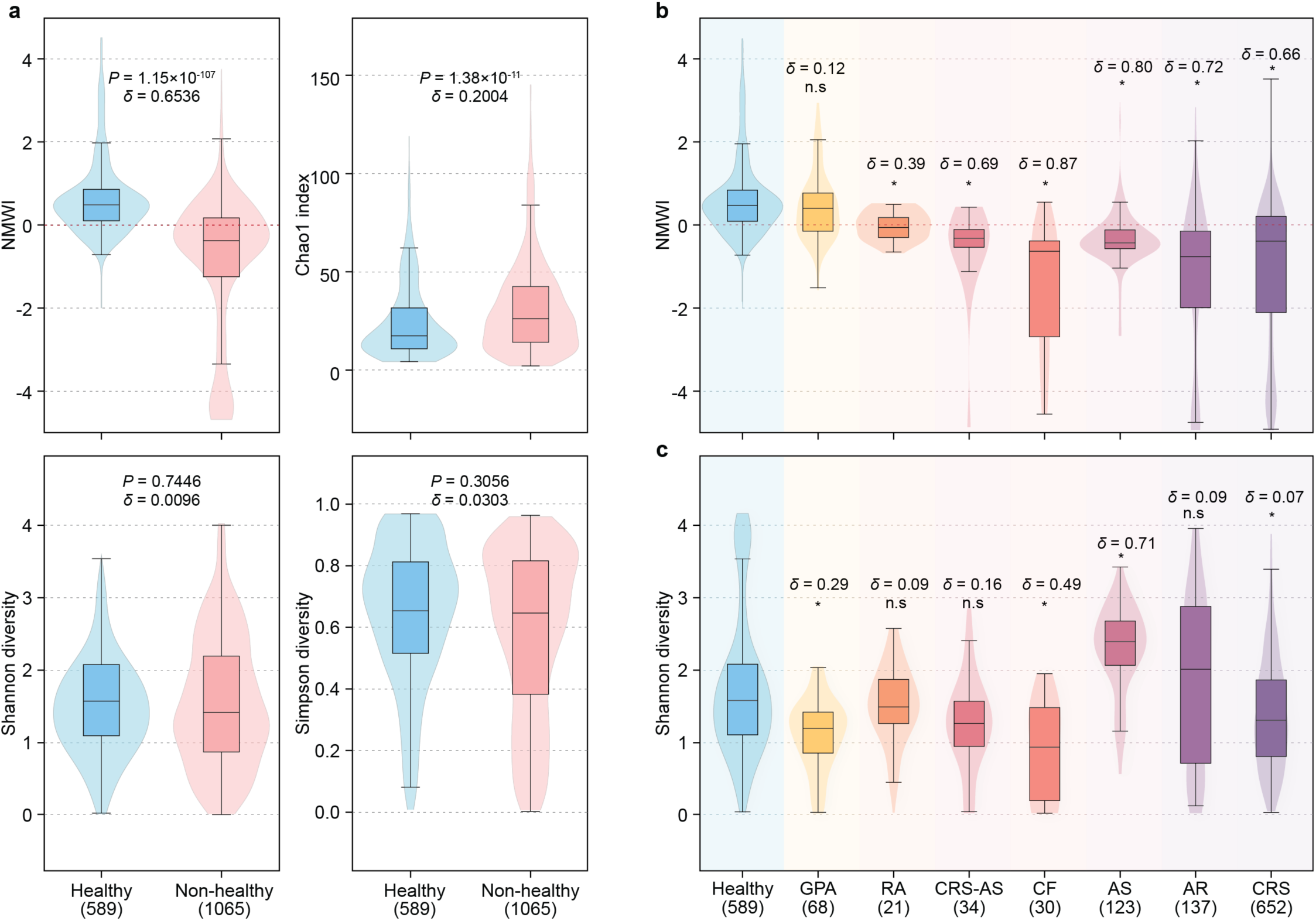
The NMWI outperforms conventional diversity metrics in distinguishing health from disease. **(a)** NMWI and diversity metrics (Shannon, Simpson, Chao1) compared between healthy and non-healthy samples (two-sided Mann–Whitney *U* tests; |Cliff’s *δ*|). Distributions of NMWI scores **(b)** and Shannon diversity **(c)** across healthy samples and individual disease cohorts (two-sided Mann–Whitney *U* tests; |Cliff’s *δ*|). Significance is denoted by * (*P* < 0.01) and n.s. (not significant).

### Predictive accuracy and generalizability of the NMWI

We evaluated predictive performance of the NMWI using complementary cross-validation and leave-out frameworks. Under LOSO CV (see **Methods**), which estimates generalizability across independent cohorts, the NMWI achieved a mean balanced accuracy of 73.82% (**Fig. 4a**). Performance was slightly lower under 10×10 CV (73.43%) and approached the resubstitution estimate (74.37%, i.e., the upper-bound estimate obtained by evaluating the model on its own training data), indicating minimal overfitting. Discrimination followed the same pattern, with AUROC values of 0.76 (LOSO CV), 0.81 (10×10 CV), and 0.83 (resubstitution) (**Fig. 4b**).

Next, to test whether the model captured generalizable rather than disease-specific signal, we performed a leave-one-disease-out (LODO) analysis, wherein we trained the model with all samples of a given disease withheld, and then evaluated it on that unseen condition. The NMWI achieved a mean balanced accuracy of 63.92% across diseases, with performance lowest for GPA (30.88%) and RA (33.33%) and highest for CF (93.33%) and AS (82.11%) (**Fig. 4c**).

To assess generalizability across 16S rRNA gene regions, we also performed a leave-one-hypervariable-region-out analysis (**Additional File 2; Figure S4**). This analysis was motivated by PERMANOVA and permutational analysis of multivariate dispersions (PERMDISP) results showing significant differences in community composition and dispersion across hypervariable regions. In each held-out iteration, samples from one hypervariable region were excluded during model training and used for testing, allowing evaluation of model performance on a previously unseen hypervariable region. In addition, to minimize bias from sample-size imbalance, this procedure was repeated 10 times, where in each iteration, a hypervariable-region group was randomly subsampled to 97 samples. The model achieved an overall mean balanced accuracy of 68.38%, with performance ranging from 65.47% to 71.44%. Thus, although hypervariable-region choice introduces measurable technical and compositional variation, the NMWI retains moderate generalizability across heterogeneous 16S rRNA gene datasets.

### NMWI thresholding and classification performance

The NMWI is defined as the predicted log-odds from an L1-regularized logistic regression model, wherein a value of zero corresponds to a predicted probability of 0.5 (indicating no class preference) and larger absolute values reflect increasing confidence toward one class. To evaluate classification under varying levels of prediction confidence, we applied a symmetric magnitude cutoff, restricting predictions to samples with stronger evidence and excluding those below the decision boundary. As scores move further away from zero in either direction, classification becomes more accurate, reflecting higher prediction confidence. In contrast, scores nearer to zero are more uncertain and less reliable in distinguishing between health states. As shown in **Figure 4d**, using 10×10 CV, the model achieved a balanced accuracy of 73.43% when no cutoff was applied with all samples classified; increasing the cutoff to 0.5 improved the balanced accuracy to 81.42% while classifying 64.19% of samples; and a cutoff of 1.0 yielded a balanced accuracy of 91.81% while retaining only 31.71% of samples (**Fig. 4e**). The same coverage–accuracy trade-off held under resubstitution (**Fig. 4f**).

**Figure 4.**
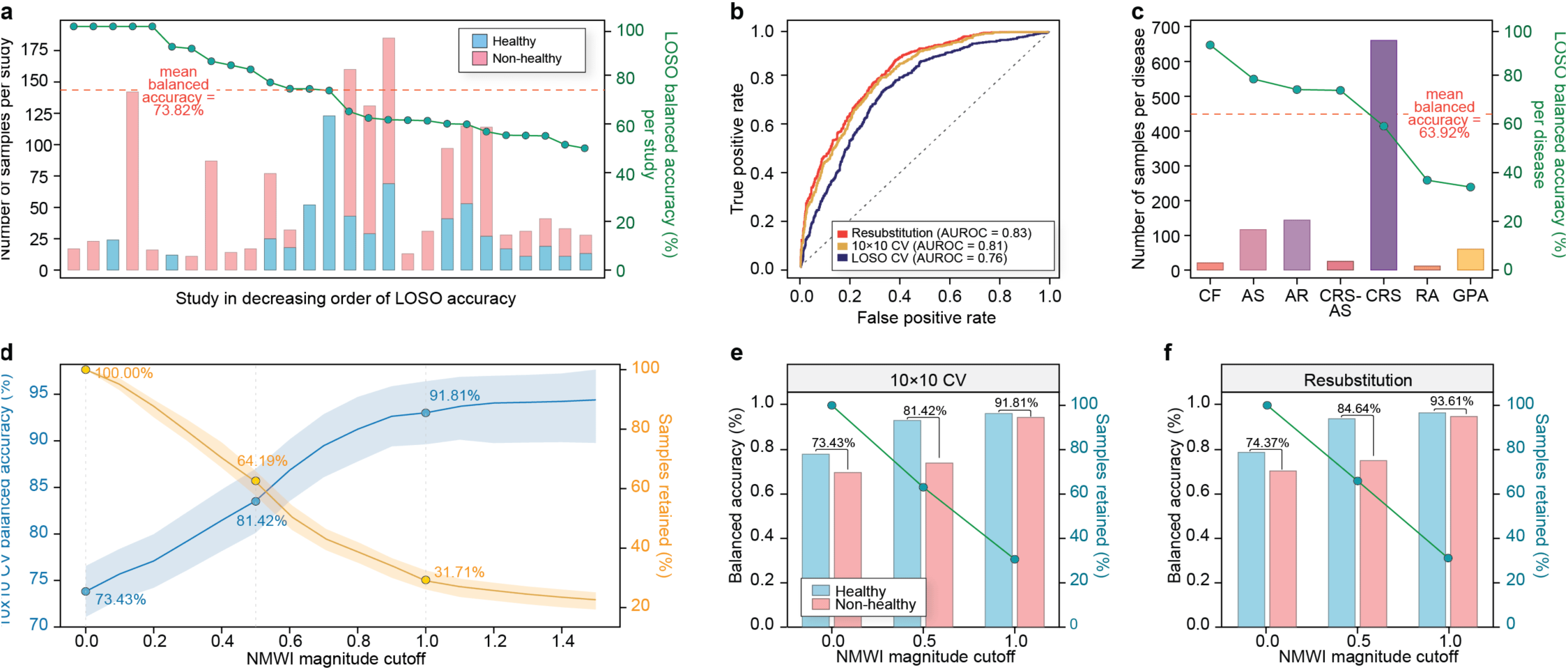
Performance and thresholding of the NMWI. **(a)** Distribution of balanced accuracies across individual studies under LOSO CV. Bars represent the number of healthy (blue) and non-healthy (red) samples per cohort, while points indicate cohort-specific balanced accuracy (green line). The dashed line denotes the cross-cohort mean (73.82%). **(b)** Receiver operating characteristic curves for classification performance in distinguishing healthy and non-healthy phenotypes under resubstitution (AUROC = 0.83), 10×10 CV (AUROC = 0.81), and LOSO CV (AUROC = 0.76). **(c)** LODO analysis of generalizability across disease conditions. **(d)** Increasing the magnitude cutoff improves classification performance, while reducing the proportion of samples retained. Balanced accuracies from 10×10 CV is shown in blue (±SD shading), with the percentage of retained samples in yellow (±SD shading). Balanced accuracies at selected magnitude cutoffs (0, 0.5, 1) under (**e**) 10×10 CV and (**f**) resubstitution. Bars show group performance; green lines indicate retained samples.

### External validation of NMWI across independent cohorts

Finally, we evaluated the generalizability of the NMWI on held-out validation datasets comprising 350 samples (109 healthy and 241 non-healthy) from 11 independent healthy and disease-specific studies. None of these samples were used at any stage of model construction, as they represented either acute conditions, consisted of only a single study per disease, or did not meet the minimum sample size requirements, and were therefore reserved for validation. With one exception, each validation study was wholly independent of the training data; Shi *et al*. [102] contributed distinct, non-overlapping disease arms to each set, with its chronic rhinosinusitis samples used in training and its anatomically and phenotypically separate NK/T-cell lymphoma samples reserved for validation. No individual sample appeared in both sets.

Encouragingly, the NMWI remained significantly greater in healthy samples compared to non-healthy samples (Mann–Whitney *U* test, *P* =1.20× 10^−17^; |Cliff’s *δ*| = 0.57; **Fig. 5a**), with a balanced accuracy of 71.49%. In contrast, Shannon diversity did not differ significantly between healthy and non-healthy samples (Mann–Whitney *U* test, *P* = 0.058; |Cliff’s *δ*| = 0.12; **Fig. 5b**), indicating only slightly lower diversity in non-healthy samples and substantial overlap between groups. Similar results were observed for Chao1 index (**Additional file 2: Figure S6**; Mann–Whitney *U* test, *P* = 0.46; |Cliff’s *δ*| = 0.05), indicating no meaningful difference between groups and negligible effect size.

At the level of individual studies, four of six healthy cohorts (or disease studies’ healthy-sample subsets) had median NMWI values above zero, yet all six differed significantly from the pooled non-healthy samples; conversely, every non-healthy cohort had a median below zero and differed significantly from the pooled healthy samples. Even where healthy-cohort medians fell below zero (as in Wu *et al*. [116] and Rhoades *et al*. [115]), they remained significantly distinct from their corresponding non-healthy samples (Wu *et al*.: *P* = 1.45 × 10^−9^, |Cliff’s *δ*| = 0.99; Rhoades *et al*.: *P* = 0.029, |Cliff’s *δ*| = 0.30). Shannon diversity and Chao1, meanwhile, did not differ consistently across cohorts (**Fig. 5b; Additional file 2: Figure S6),** with most comparisons non-significant and variable in direction. Together, these results demonstrate that the NMWI maintains robust, consistent discrimination of nasal microbiome health states across fully independent datasets, outperforming conventional diversity metrics in external validation.

**Figure 5.**
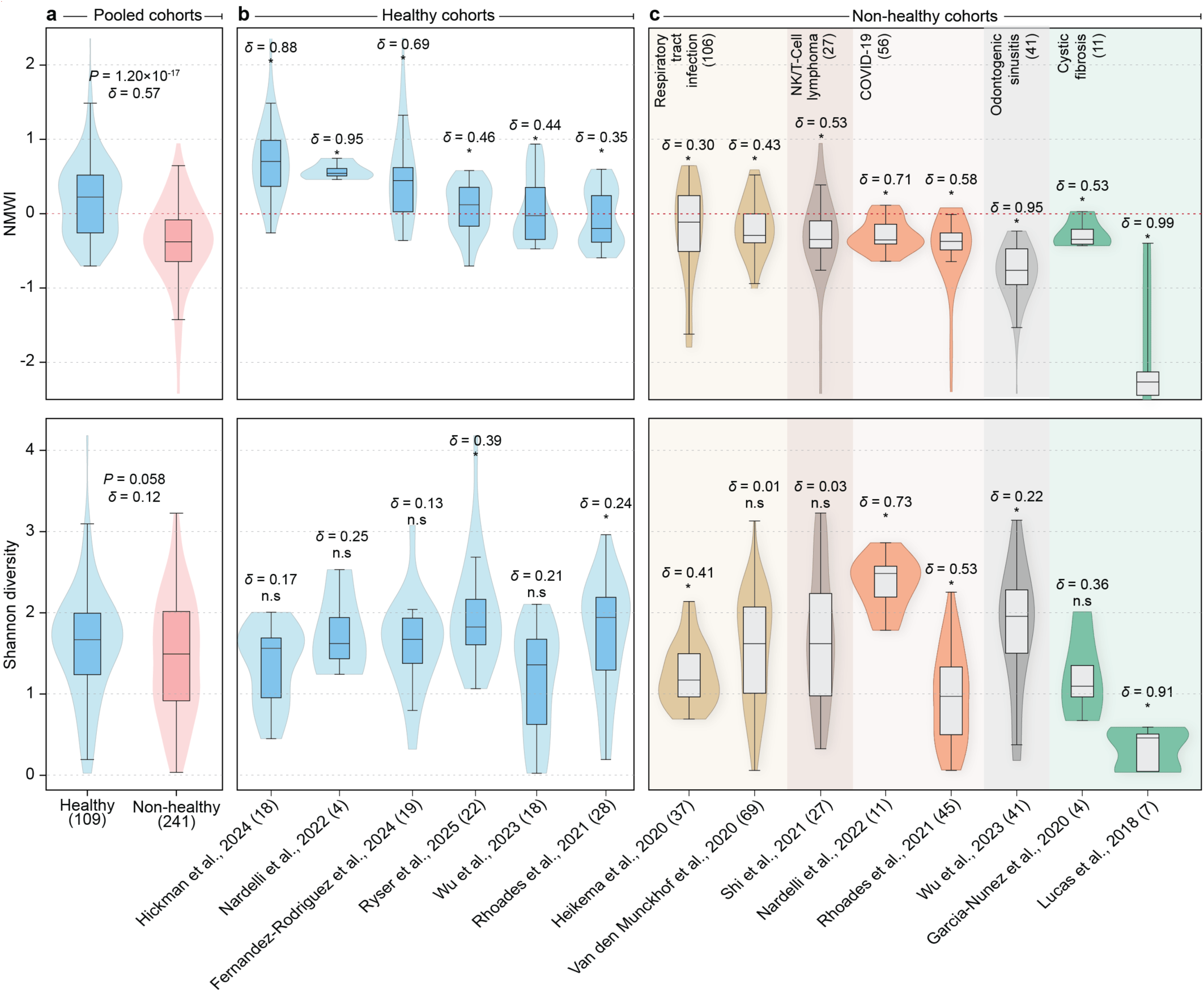
Validation of the NMWI across independent datasets and disease phenotypes. Top panels display NMWI score distributions, and bottom panels show Shannon diversity in the same validation cohort for comparison. **(a)** Pooled healthy vs. non-healthy samples, demonstrating significant separation in NMWI scores (Mann–Whitney *U* test, *P* = 1.20×10^−17^; |Cliff’s *δ*| = 0.57), whereas Shannon diversity did not differ significantly (*P* = 0.058, |*δ*| = 0.12). **(b)** Distributions across independent healthy cohorts. Asterisks indicate statistically significant differences relative to pooled non-healthy samples (Mann–Whitney *U* test, *P* < 0.05). **(c)** Study-specific distributions across non-healthy cohorts. Asterisks denote significant differences relative to pooled healthy samples (Mann–Whitney *U* test, *P* < 0.05).

## Discussion

The Nasal Microbiome Wellness Index (NMWI) represents a meaningful advance in translational nasal microbiome research: a disease-agnostic, standardized, and interpretable index of nasal microbiome health status derived from a large, uniformly reprocessed multi-study dataset. (To our knowledge, no equivalent framework existed for the nasal cavity prior to this work.) By training a LASSO logistic regression model on 1654 healthy (disease free) and non-healthy (disease harboring) nasal microbiome profiles—pooled across 27 publicly available 16S rRNA gene datasets spanning seven chronic disease conditions and four continents—we identified a sparse set of microbial features and estimated their coefficients, which together compute a single score: the predicted log-odds that a profile resembles a healthy nasal state. A key design principle was to restrict model training to stable chronic disease cohorts, avoiding the transient and condition-specific perturbations associated with acute illness, and thereby prioritizing stable, ecologically meaningful signals over short-term host responses. The result is an index that, across multiple validation frameworks, robustly and consistently discriminates healthy from non-healthy nasal microbiome states—a capability that conventional α-diversity metrics consistently failed to achieve in the same datasets.

Across cross-validation and leave-out frameworks, the NMWI achieved a mean balanced accuracy of 73.82% under leave-one-study-out cross-validation (LOSO CV) and 73.43% under 10×10 CV, both approaching the resubstitution estimate (74.37%). The near-identical performance observed under LOSO CV and 10×10 CV indicates that the model is robust to study-level heterogeneity and does not rely heavily on study-specific compositional signatures. This finding is notable given the substantial variability in cohort demographics, sampling protocols, and geographic origins represented across studies [126,127]. Despite these sources of heterogeneity, the LOSO CV mean balanced accuracy remained above chance for all but the most biologically challenging cohorts, and the overall AUROC pattern (0.76 LOSO CV vs. 0.83 resubstitution) is consistent with a model that captures genuine cross-study signal rather than overfitted study-specific patterns. Critically, the model maintained a balanced accuracy of 71.49% on external validation samples excluded from all stages of model development, providing strong evidence of generalizability to unseen data.

The inadequacy of α-diversity metrics as markers of nasal microbiome health was not merely an expectation based on previous findings [68,69], but a consistent empirical finding across every dataset examined in our study. Among the diversity metrics evaluated, only Chao1 differed significantly between pooled healthy and non-healthy groups. However, the effect size remained small (|Cliff’s *δ*| ≤ 0.20). The remaining diversity metrics did not reach statistical significance, and their distributions were almost entirely overlapping. More strikingly, the diversity metrics showed directional inconsistency across disease conditions—some diseases showed higher diversity than healthy controls and others lower, with Chao1 and Shannon at times pointing in opposite directions for the same condition. These inconsistencies underscore a fundamental interpretive problem: in a body site characterized by low-diversity communities dominated by a limited set of niche-adapted taxa [10,11], diversity metrics are particularly prone to reflecting stochastic compositional fluctuations rather than meaningful health-associated shifts. The NMWI, by contrast, produced consistent negative shifts (relative to healthy controls) across all disease conditions examined (|Cliff’s *δ*| = 0.12–0.87), a pattern that held in the independent external validation cohort (|Cliff’s *δ*| = 0.30–0.99). These results support the argument for moving beyond diversity metrics toward integrative, ecologically informed indices as the primary tool for microbiome health assessment [128].

The leave-one-disease-out (LODO) analysis further strengthens the case for NMWI’s disease-agnostic design. By iteratively retraining the model on all diseases except one and evaluating performance on the held-out condition, we demonstrated that the NMWI captures microbial health signatures that transfer across disease contexts, not merely within them. Mean balanced accuracy across LODO iterations was 63.92%, with performance varying substantially across conditions—ranging from 30.88% in granulomatosis with polyangiitis (GPA) to 93.33% in cystic fibrosis (CF). These differences are biologically informative rather than simply reflecting model limitations. Reduced discrimination in GPA may be attributable to the species-level specificity of its microbiome alterations: prior studies have shown that GPA-associated nasal dysbiosis is driven primarily by opposing shifts within the genus *Staphylococcus*—specifically an increase in *S. aureus* and depletion of *S. epidermidis* [103,104,129]—rather than broad community restructuring. Because genus-level 16S rRNA gene data cannot resolve such intrageneric dynamics, NMWI sensitivity is inherently limited in this context. Similarly, the reduced LODO accuracy for RA (33.33%) is consistent with the modest, heterogeneous nasal microbiome alterations reported for this systemic autoimmune condition [108]. In contrast, the high LODO accuracy for CF (93.33%) and asthma (AS, 82.11%) aligns with the pronounced, well-documented, microbial shifts in these conditions, including expansion of gram-negative taxa such as *Pseudomonas* in CF airways [91,92].

The 24 taxa retained by the LASSO model are biologically consistent with the established ecology of the healthy adult nasal microbiome and offer interpretive clarity that neither diversity metrics nor differential abundance lists provide alone. Positive contributors (e.g., *Corynebacterium*, *Cutibacterium*) reflect the well-characterized core of stable, healthy nasal communities [58,67,130–132]. *Corynebacterium accolens,* for example, releases antimicrobial fatty acids that inhibit *Streptococcus* colonization [130,133], while *Cutibacterium* species have been associated with a balanced nasal immune environment and protective mucosal responses [123]. In contrast, the negative contributors (e.g., *Pseudomonas*, *Escherichia–Shigella*, *Haemophilus*) are consistent with previous reports of their association with dysbiotic nasal states, chronic upper-airway disease, and gram-negative-dominated communities [44,75,98,100,121]. Notably, strictly anaerobic taxa, including members of the *Lachnospiraceae* family, *Fusobacterium* spp, and *Prevotella* spp, were also identified as contributors to lower NMWI scores. This finding is consistent with the altered ecological conditions observed in inflammatory sinonasal diseases including reduced oxygen tension and increased acidity, which favors the expansion of anaerobes [26,134,135]. Despite being often considered gut-associated anaerobes, *Lachnospiraceae* are also part of the microbial landscape of nasal, oropharyngeal, and sinus-associated niches [5,26,105,135–138]. Their metabolic capacity for butyric acid formation has been associated with altered olfactory function and localized inflammatory responses [26,105,135,138]. Importantly, these associations reflect the joint predictive contribution of multiple taxa operating as a community-level signal, not independent effects—a distinction that is critical in the sparse, low-diversity ecology of the nasal cavity, where a handful of dominant genera account for most of the community and the relative balance among taxa carries more information than the abundance of any single organism. The NMWI formalizes this intuition by weighting each taxon’s contribution according to its LASSO-derived coefficient, producing a score that reflects the community’s configuration as a whole.

One of the most technically distinctive and consequential aspects of the NMWI is that it was derived entirely from 16S rRNA gene amplicon sequencing data spanning five hypervariable regions (V1–V2, V1–V3, V1–V4, V3–V4, and V4) across 27 independently published studies. (Notably, the model then generalized to an *unseen* sixth region [V5–V6] in external validation.) This stands in direct contrast to the GMWI and GMWI2, which were trained exclusively on shotgun metagenomics data. The challenge inherent in our approach is substantial: different hypervariable regions carry different taxonomic biases, resolution, and primer efficiencies, introducing systematic variation in community composition estimates that can be analytically indistinguishable from true biological differences [65,66]. That a generalizable, cross-cohort health signal emerges despite this methodological heterogeneity is itself a notable finding—it demonstrates that the nasal microbiome carries community-level health signatures robust enough to be detected across methodological variation.

This multi-region 16S approach also confers important practical advantages. Shotgun metagenomics, while offering superior taxonomic and functional resolution, remains more resource-intensive, requires greater sequencing depth, and is unavailable for the vast majority of clinical and epidemiological nasal microbiome datasets in the public domain. In contrast, 16S rRNA gene sequencing is the dominant methodology in existing nasal microbiome research, and so the NMWI is directly applicable to the majority of published datasets without re-sequencing—dramatically broadening the scope of reanalysis studies, retrospective cohort applications, and longitudinal investigations that can immediately leverage it. Furthermore, the demonstration that a LASSO model trained on multi-region 16S data generalizes to held-out studies suggests that genus-level taxonomic profiles—the resolution level at which most 16S data is confidently interpretable [139]—capture sufficient community structure for health status classification. This finding extends beyond the NMWI itself, providing empirical support for harmonized genus-level 16S data in multi-study machine learning applications in the nasal and, potentially, other body-site microbiomes.

The NMWI possesses several features that make it well suited for translational applications in respiratory precision medicine, particularly of the upper airway. Nasal sampling is non-invasive, inexpensive, and repeatable, enabling longitudinal health monitoring with high adherence [60]. The continuous nature of the NMWI score supports nuanced, patient-level interpretation: rather than forcing binary healthy/non-healthy assignments, clinicians or researchers can treat the magnitude as a measure of prediction confidence, and the magnitude-thresholding framework introduced here allows users to tune the trade-off between coverage and certainty for a given application. At a cutoff of 1.0 in 10×10 CV, 91.81% balanced accuracy was achieved among the 31.71% of samples classified—a performance profile analogous to the “reject option” framework used in GMWI2 [72] and directly relevant to clinical screening, where high-confidence calls are preferable to universal but uncertain assignments. Longitudinal applications are a particularly compelling future direction: just as GMWI2 has been applied to track gut microbiome health through dietary interventions, antibiotic exposure, and fecal microbiota transplantation [72], the NMWI is conceptually suited to track nasal microbiome trajectories in response to corticosteroid therapy [140], endoscopic surgery for chronic rhinosinusitis [141], or biologic therapy such as dupilumab [113]. Lastly, because the nasal cavity is the primary interface between the host and the external environment, it is uniquely positioned to capture the microbial consequences of environmental exposures (e.g., air pollution [142,143], occupational exposures [144], seasonal allergens [145]) through non-invasive sampling. The NMWI therefore offers a practical tool not only for clinical monitoring but for environmental health research, enabling standardized, quantitative assessments of how ambient exposures shape and perturb nasal microbiome health at the population level.

Several limitations warrant consideration. First, the pooled design integrates studies differing in sampling site, primer region, extraction protocol, sequencing platform, and metadata completeness. Although standardized reprocessing substantially reduces technical variability, residual batch effects cannot be fully eliminated, and their contribution to cross-study heterogeneity in NMWI performance cannot be fully disentangled from genuine biological variation. Publicly available datasets also represent a potentially skewed subset of the broader population, shaped by study-specific recruitment criteria and protocols, which may limit generalizability to the general public. Second, restriction to genus-level resolution—a consequence of 16S rRNA gene data—limits sensitivity to intrageneric dynamics. This is particularly consequential for *Staphylococcus*, where opposing species-level shifts (*S. aureus* vs. *S. epidermidis*) may cancel at the genus level and reduce discriminability in conditions such as GPA [103,104,129]. This limitation is clinically salient: *S. aureus* nasal colonization is among the most consequential microbial features assessed in routine upper-airway care, and resolving it would be of substantial clinical value. Strain- and species-resolved approaches—whether targeted assays or shotgun metagenomics—are therefore a high priority for future iterations of the index. Third, the cross-sectional design of most included studies precludes causal inference. The NMWI should be interpreted as an associative rather than mechanistic marker of health status; our data cannot establish whether dysbiotic configurations contribute to disease or instead arise as a consequence of impaired host health, and this directionality has direct implications for therapeutic actionability. Distinguishing these possibilities—and determining whether modulating the nasal microbiome can shift the NMWI and, in turn, clinical outcomes—will require longitudinal and interventional studies. Fourth, host covariates (e.g., sex, age, BMI, smoking status, medications) were inconsistently reported across studies and could not be systematically incorporated, leaving open whether NMWI performance varies across demographic subgroups. Fifth, the current model is trained on 16S rRNA gene data and is not directly applicable to shotgun metagenomic profiles without retraining, limiting cross-platform comparability with gut microbiome wellness indices.

A further consideration concerns the scope of the non-healthy group. With the exception of RA, the chronic conditions modeled here involve direct nasal or respiratory pathology, an appropriate focus for an index designed to assess upper-airway health. It remains an open and intriguing question whether the nasal microbiome also reflects systemic health states without overt airway involvement—such as cardiometabolic disease, obesity, or malignancy—as has increasingly been described for the gut microbiome. Clinical experience suggests this possibility is not far-fetched: *S. aureus* nasal colonization, for example, does not invariably accompany worse localized disease, hinting that nasal microbial state may at times index broader host physiology rather than strictly local airway health. Extending the NMWI framework to systemic disease cohorts would help delineate whether nasal microbiome health is a local or a more global marker, and represents a natural direction for future work.

## Conclusions

By reducing complex nasal microbial profiles to a single interpretable score, the NMWI moves the field beyond descriptive taxonomic comparison toward a quantitative, standardized framework suited to individual-level health assessment, cross-study benchmarking, and the tracking of microbiome responses to clinical intervention and environmental exposure. As an open-source, extensible resource, our index establishes both a practical tool and a methodological precedent for quantitative microbiome-based phenotyping in respiratory precision medicine. More broadly, by rendering nasal microbiome health measurable and repeatable, the NMWI supports a longer-term shift in upper-airway care—from reactive assessment after symptoms emerge toward proactive, non-invasive monitoring and earlier risk stratification.

## Declarations

### Ethics approval and consent to participate

Not applicable.

### Consent for publication

Not applicable.

### Availability of data and materials

All raw 16S rRNA gene sequencing data (FASTQ files) used in this study are publicly available through the NCBI Sequence Read Archive (SRA). The training and validation datasets included in the analysis are summarized in **Table 1**. Full metadata and SRA accession numbers for the training datasets are provided in **Additional file 1: Tables S1**, **S2,** respectively, and those for the validation datasets are provided in **Additional file 1: Tables S3**, **S4**, respectively. Processed datasets and analysis code generated during this study are available in the project GitHub repository: https://github.com/mabarrera12/NMWI/.

### Authors’ contributions

M.A.B.-S., V.K.G., V.L.K., and J.S. developed the study idea and designed all analytical methodologies. M.A.B.-S. performed the computational experiments. All authors (M.A.B.-S., V.K.G., X.Z., A.L.K., C.Y.Z., G.S., L.V.K., E.K.O., V.L.K., and J.S.) critically reviewed and discussed the data. M.A.B.-S. and J.S. wrote the manuscript, with editorial contributions from other authors. All authors approved the final manuscript.

