## Supplementary Figures for "A Disease-Agnostic Nasal Microbiome Wellness Index for Standardized Assessment of Upper-Airway Respiratory Health"

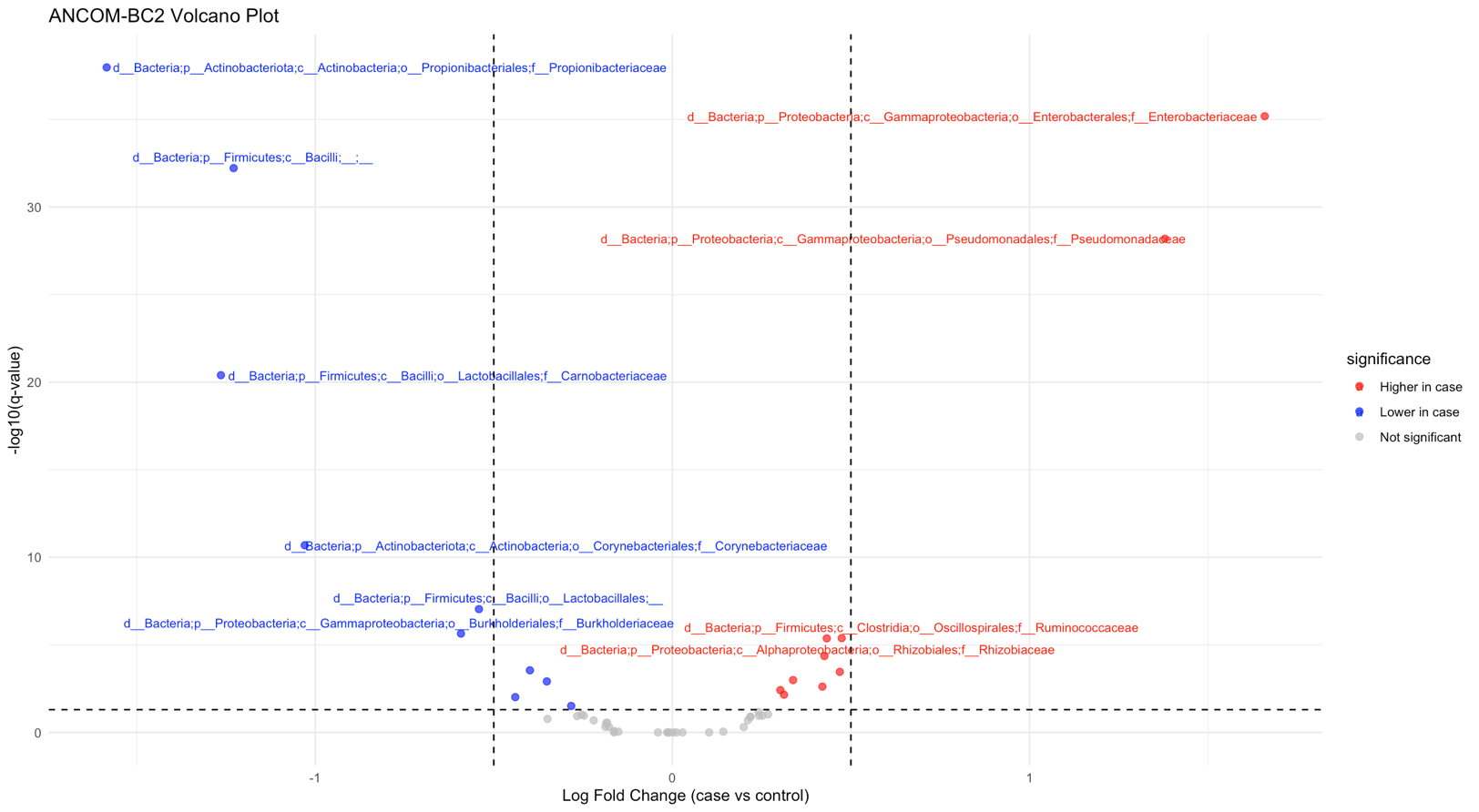

**Figure S1: Differential abundance analysis at the family-level between healthy and non-healthy groups.** Volcano plot of ANCOM-BC2 family-level differential abundance results. Each point represents a bacterial family; the x-axis shows natural log fold change (LFC) (non-healthy vs. healthy), and the y-axis shows −ln(adjusted *P*) (Holm-adjusted). Positive values indicate higher abundance in non-healthy samples, negative values in healthy samples. Dashed lines mark (LFC) and significance thresholds (adj *P* = 0.05). Red points are significantly higher in non-healthy, blue in healthy, and grey are non-significant.

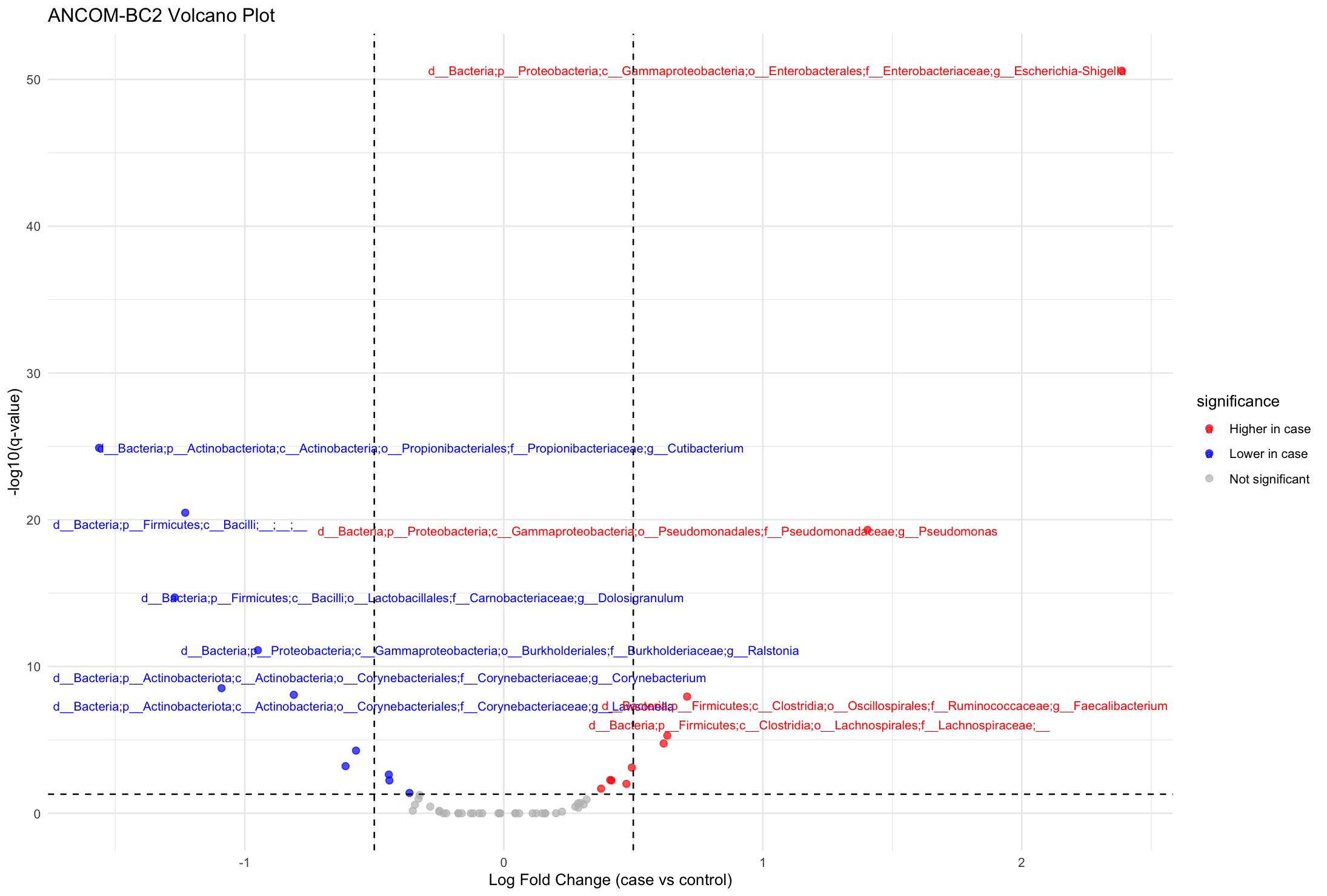

**Figure S2: Differential abundance analysis at the genus-level between healthy and non-healthy groups.** Volcano plot of ANCOM-BC2 genus-level differential abundance results. Each point represents a bacterial genus; the x-axis shows natural log fold change (LFC) (non-healthy vs. healthy), and the y-axis shows −ln(adjusted *P*) (Holm-adjusted). Positive values indicate higher abundance in non-healthy samples, negative values in healthy samples. Dashed lines mark (LFC) and significance thresholds (adj *P* = 0.05). Red points are significantly higher in non-healthy, blue in healthy, and grey are non-significant.

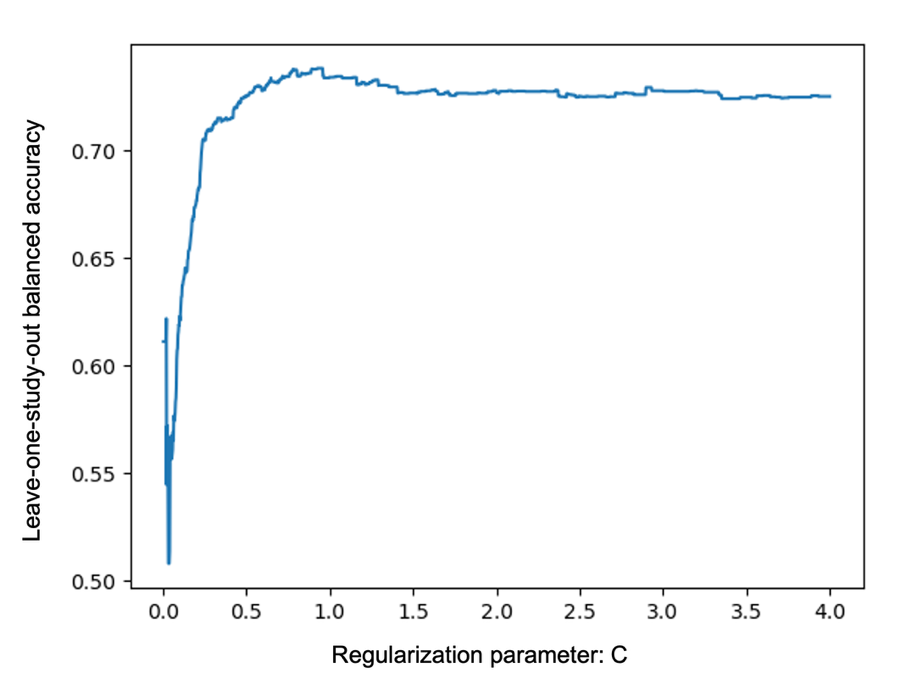

**Figure S3**: **Hyperparameter optimization of the L1-regularized logistic regression model.** Leave-one-study-out cross-validation (LOSO CV) performance across a grid of regularization parameter values (*C*). The x-axis shows *C* (inverse regularization strength), and the y-axis shows mean balanced accuracy within LOSO CV. The optimal *C* was selected as the value yielding the highest mean balanced accuracy.

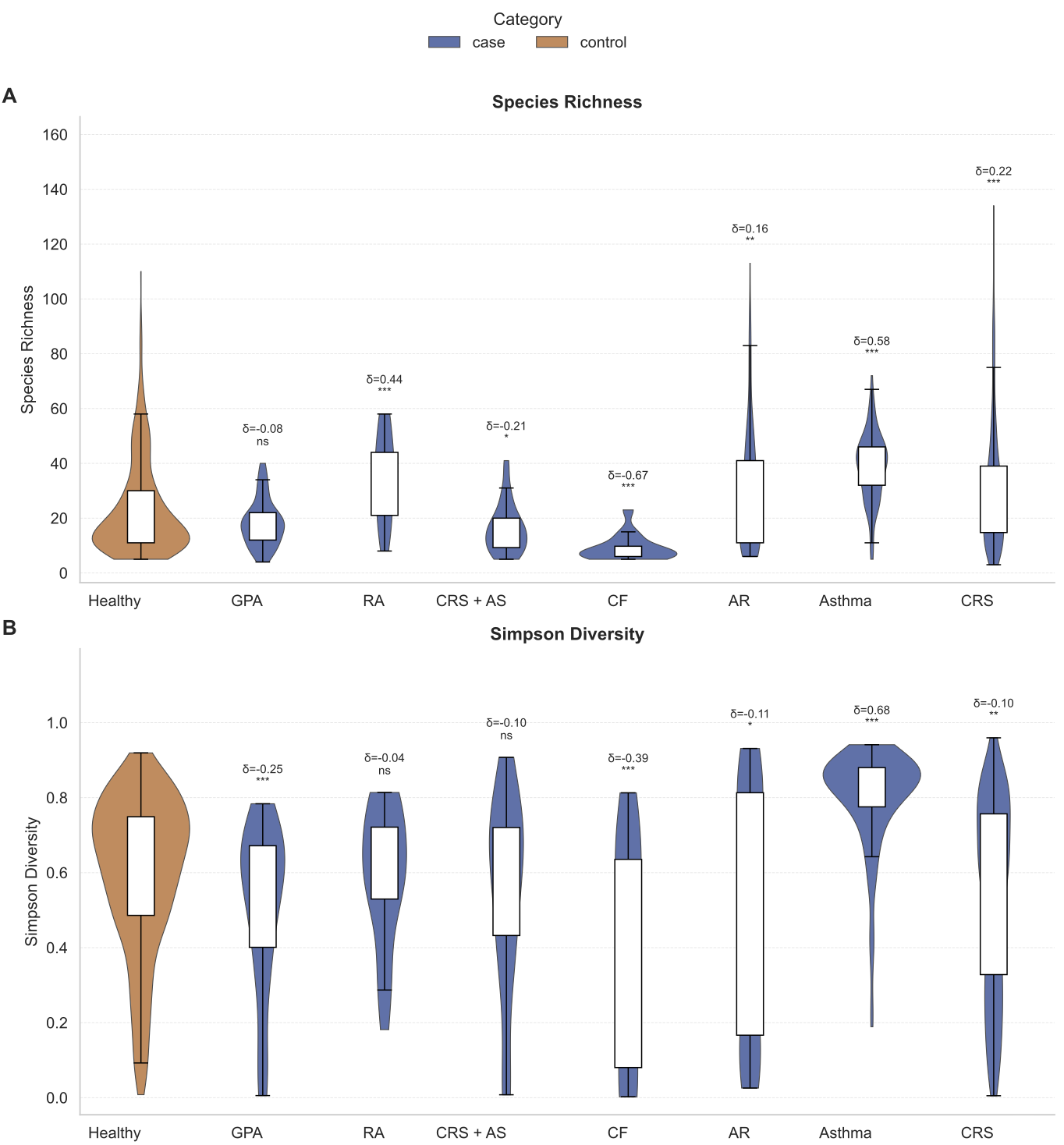

**Figure S4: Alpha-diversity across healthy and disease groups. A)** Species richness (Chao1) across healthy controls and disease groups (GPA, RA, CRS-AS, CF, asthma, AR, CRS). **B)** Simpson diversity index for the same groups. Violin plots depict the distribution of values within each group, with embedded boxplots indicating the median and interquartile range. Healthy controls (orange) are shown alongside case groups (blue). Cliff’s delta (*δ*) effect sizes and Mann–Whitney *U* test significance are displayed above each disease group relative to healthy controls. Positive *δ* values indicate higher diversity in disease cohorts, whereas negative values indicate higher diversity in healthy controls. Statistical significance is denoted as *P* < 0.05 (*), *P* < 0.01 (**), and *P* < 0.001(***); “ns” indicates a non-significant difference.

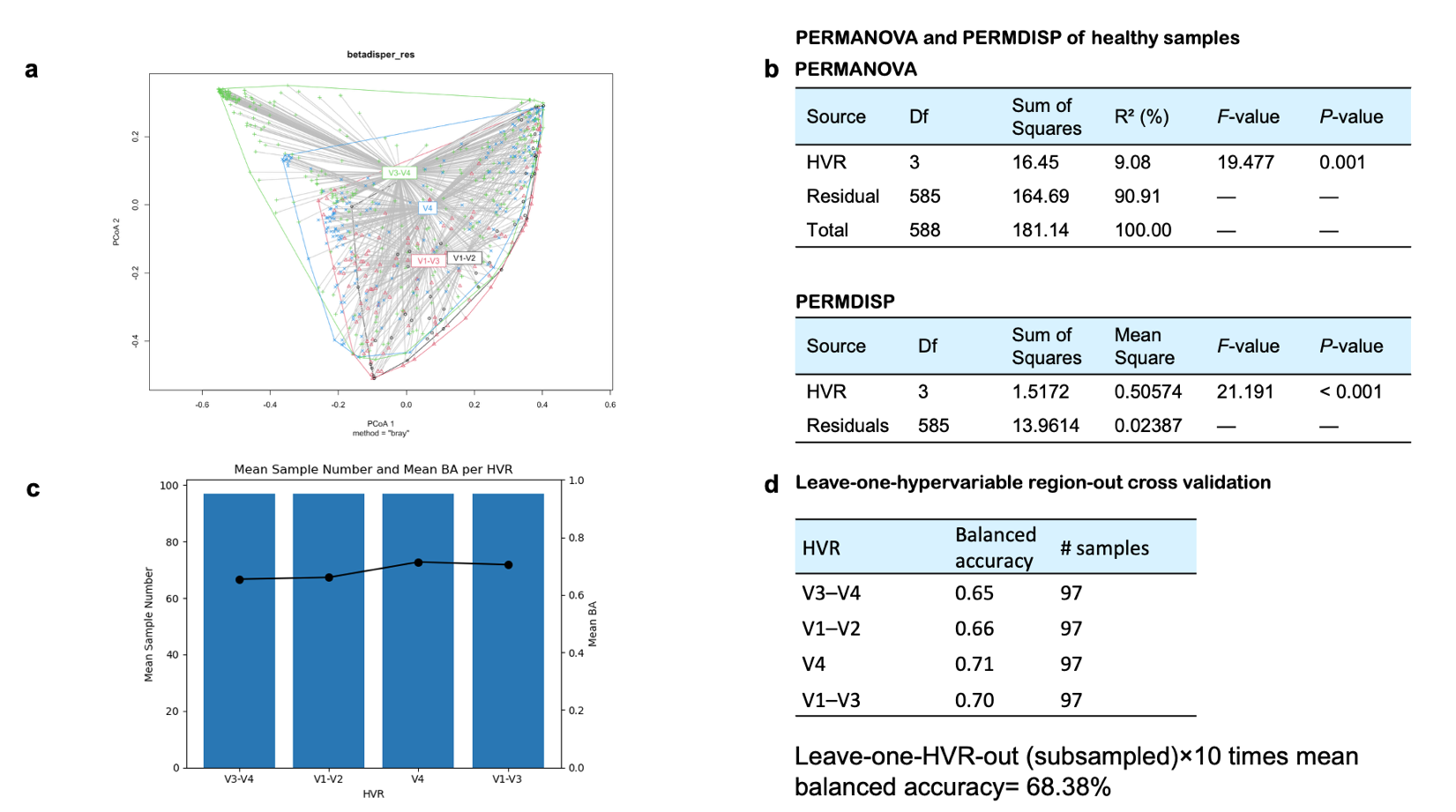

**Figure S5: Influence of 16S rRNA gene hypervariable region on NMWI classification performance.** Analyses were performed in R using the *vegan* package for Bray–Curtis dissimilarity, PERMANOVA, and PERMDISP. **(a)** PCoA of Bray–Curtis dissimilarity showing clustering of healthy nasal microbiome samples by amplified hypervariable region (HVR). **(b)** PERMANOVA and PERMDISP results assessing differences in community composition and dispersion across HVR groups. Significant PERMANOVA results should be interpreted in the context of the accompanying PERMDISP analysis, which also detected significant differences in dispersion among HVR groups, indicating that both centroid separation and within-group variability contributed to the observed compositional differences. **(c)** Sample number and balanced accuracy per HVR after subsampling, showing that NMWI performance was evaluated under balanced sample-size conditions across HVR groups. **(d)** Leave-one-HVR-out validation of NMWI performance. In each analysis, one HVR group was withheld from model training and used as the test set. For each held-out HVR, samples were subsampled 10 times to 97 samples to reduce sample-size imbalance, and balanced accuracy was averaged across subsampling iterations. Despite measurable HVR-associated differences in community composition and dispersion, the NMWI retained above-chance performance across held-out HVRs, with an overall mean balanced accuracy of 68.38% and HVR-specific values ranging from 65% to 71%. These results indicate that HVR choice introduces technical and compositional variation but does not fully account for NMWI discrimination.

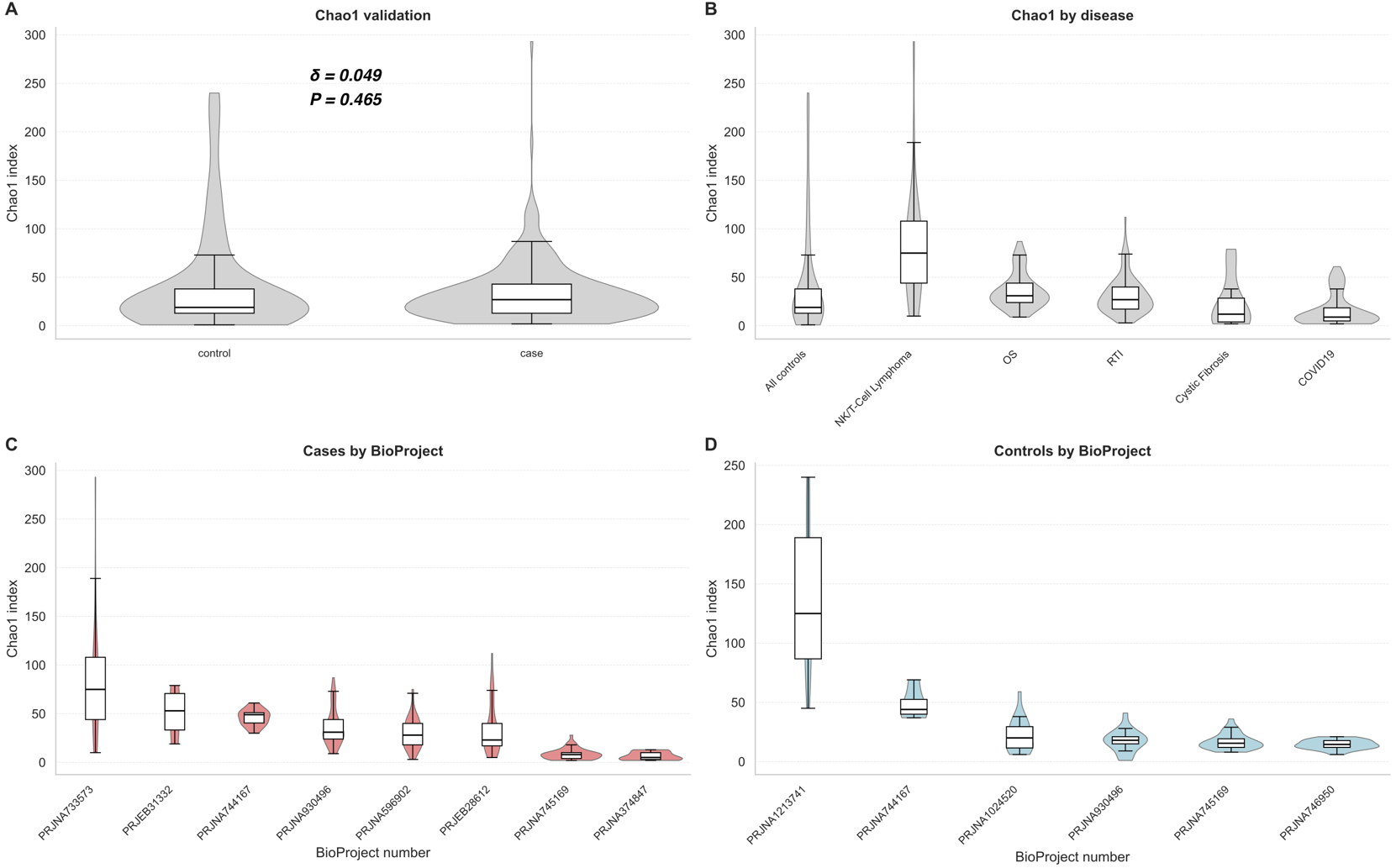

**Figure S6: Chao1 richness across external validation cohort.** **A)** Overall comparison of Chao1 richness between pooled healthy controls and non-healthy samples. Cliff’s delta (*δ*) effect sizes and Wilcoxon rank-sum test significance (*P*) are displayed relative to healthy controls. **B)** Chao1 richness distributions for individual disease cohorts **C)** Chao1 richness stratified by BioProject among non-healthy samples. **D)** Chao1 richness stratified by BioProject among healthy controls. Violin plots represent the distribution of values within each group, and embedded boxplots indicate the median and interquartile range.
